## Supplementary Material for "multimedia: Multimodal Mediation Analysis of Microbiome Data"

October 3, 2024

Table 1: National Cancer Institute Quick Food Scan (Part). Instructions: Think about your eating habits over the past 12 months. About how often did you eat or drink each of the following foods? Remember breakfast, lunch, dinner, snacks, and eating out. Blacken in only one bubble for each food.

| Type of Food | Never | Less than Once Per Month | 1-3 Times Per Month | 1-2 Times Per Week | 3-4 Times Per Week | 5-6 Times Per Week | 1 Time Per Day | 2 or More Times Per Day |
|---|---|---|---|---|---|---|---|---|
| Cold Cereal | ○ | ○ | ○ | ○ | ○ | ○ | ○ | ○ |
| Fruit (not juices) | ○ | ○ | ○ | ○ | ○ | ○ | ○ | ○ |

Table 2: PROMIS-43 Profile v2.0 (Part). Instructions: respond to each question or statement by marking one box per row.

| | | | | | |
|---|---|---|---|---|---|
| **Fatigue** | | | | | |
| **During the past 7 days ...** | Not at all | A little bit | Somewhat | Quite a bit | Very much |
| I feel fatigued | ☐ | ☐ | ☐ | ☐ | ☐ |
| I have trouble starting things because I am tired | ☐ | ☐ | ☐ | ☐ | ☐ |
| **In the past 7 days ...** | | | | | |
| How run-down did you feel on average | ☐ | ☐ | ☐ | ☐ | ☐ |
| How fatigued were you on average | ☐ | ☐ | ☐ | ☐ | ☐ |
| How much were you bothered by your fatigue on average | ☐ | ☐ | ☐ | ☐ | ☐ |
| To what degree did your fatigue interfere with your physical functioning | ☐ | ☐ | ☐ | ☐ | ☐ |
| **Sleep Disturbance** | | | | | |
| **In the past 7 days ...** | Very Poor | Poor | Fair | Good | Very Good |
| My sleep quality was | ☐ | ☐ | ☐ | ☐ | ☐ |
| **In the past 7 days ...** | Not at all | A little bit | Somewhat | Quite a bit | Very much |
| My sleeping was refreshing | ☐ | ☐ | ☐ | ☐ | ☐ |
| I had a problem with my sleep | ☐ | ☐ | ☐ | ☐ | ☐ |
| I had difficulty falling sleep | ☐ | ☐ | ☐ | ☐ | ☐ |
| My sleep was restless | ☐ | ☐ | ☐ | ☐ | ☐ |
| I tried hard to get to sleep | ☐ | ☐ | ☐ | ☐ | ☐ |

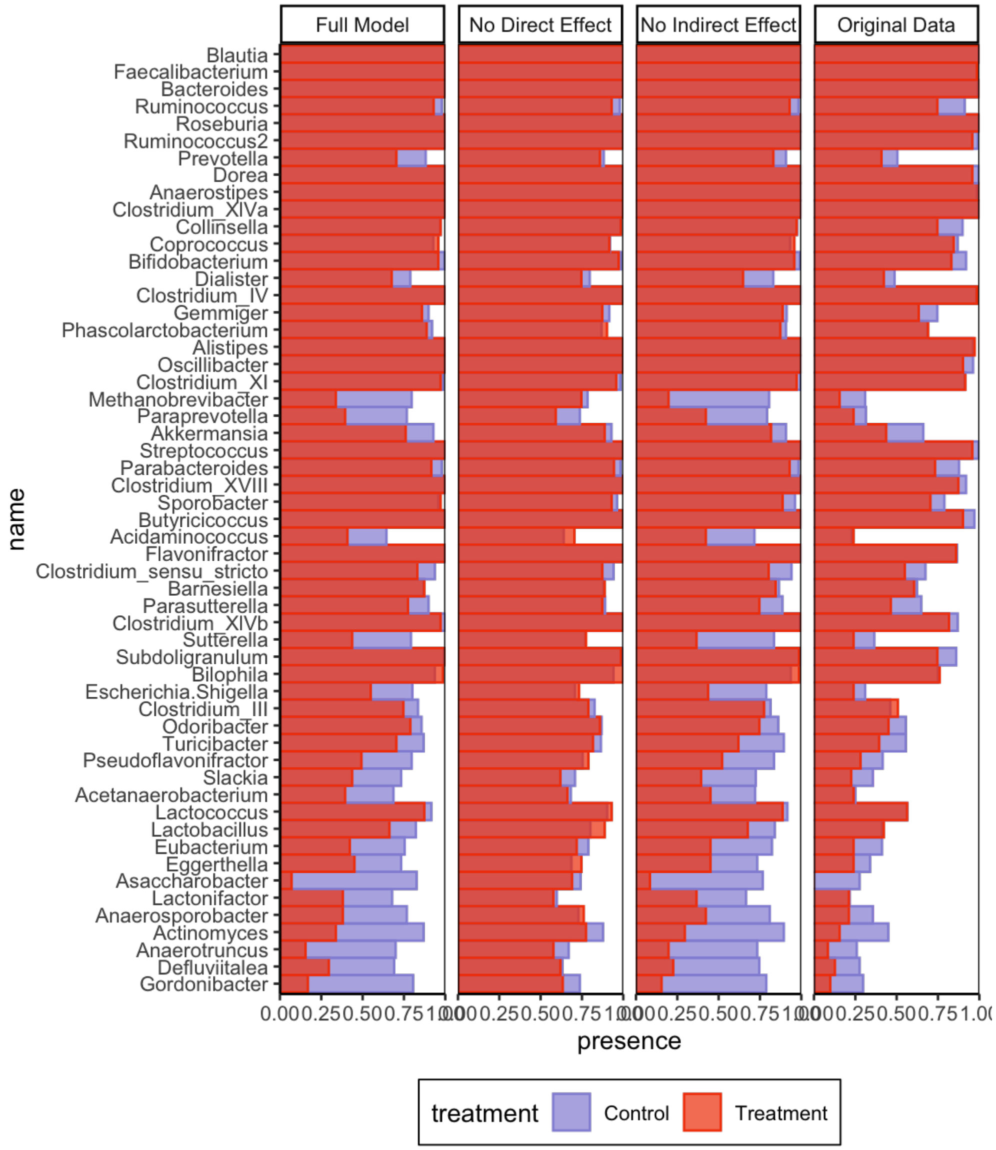


Figure 1: Simulated and real presence proportions for the logistic normal multinomial models discussed in the mindfulness case study. Taxa with inappropriately high spread in simulated proportions in Fig. **7** tend to be observed in fewer real data samples, despite having high abundance when observed. None of the synthetic datasets reflect this phenomenon, indicating a basic limit of the logistic normal multinomial model in modeling zero inflation.

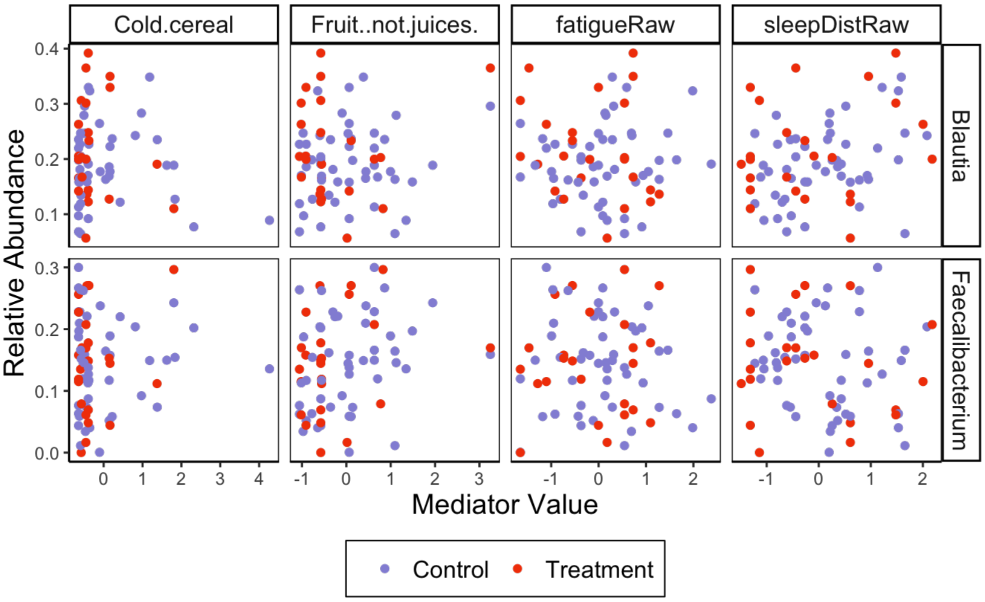


Figure 2: Associations between mediators and relative abundance for the two taxa selected by the synthetic null hypothesis testing procedure applied to overall indirect effects (Fig. **8**). Indirect effects exist when the mediator is both affected by the treatment and correlated with the response.
